# Long Term Culture of Germ-Free Zebrafish Using Gamma-Irradiated Feeds

**DOI:** 10.1101/2025.10.09.681431

**Authors:** Lydia Okyere, Angela Di Fulvio, Christopher Gaulke

**Affiliations:** Department of Pathobiology, University of Illinois Urbana-Champaign; Department of Nuclear, Plasma, and Radiological Engineering, University of Illinois Urbana-Champaign; Carl R. Woese Institute for Genomic Biology, University of Illinois Urbana-Champaign; Personalized Nutrition Initiative, University of Illinois Urbana-Champaign; Cancer Center at Illinois, University of Illinois Urbana-Champaign; National Center for Supercomputing Applications, University of Illinois Urbana-Champaign; Interdisciplinary Health Sciences Institute, University of Illinois Urbana-Champaign

**Keywords:** Zebrafish, gut, microbiome, gnotobiotic, germ-free, host-microbe interactions

## Abstract

Host associated microbiota play essential roles in regulating digestion, nutrient acquisition, immunity, and xenobiotic metabolism. Disruption of these communities is linked to numerous diseases and health defects though causal mechanisms underpinning these associations remain unclear in most cases. Gnotobiotic zebrafish provide a scalable low-cost *in vivo* model that is increasingly used to resolve causality in host-microbiota interactions. However, reliance on live diets limits the use of gnotobiotic zebrafish to early life stages where body systems and microbial communities are incompletely developed. As a result, many important host-microbiota interactions may be unable to be studied in this model system. Here we tested a simple method for long-term husbandry of gnotobiotic zebrafish using gamma-irradiated chow diets and evaluated effects on growth, gene expression, and microbial community composition. In conventionally reared animals, gamma irradiated diets did not affect growth or survival and only modestly impacted microbial community composition and diversity. In contrast, gnotobiotic zebrafish maintained on sterile irradiated diets for 55 days post fertilization were smaller, weighed less, and exhibited aberrant genes expression profiles relative to controls. These genes were enriched for pathways, related to immune response, xenobiotic metabolism, organ development, liver function, and lipid metabolism, with many expression patterns linked to the abundance of specific microbial taxa. Together, these findings establish a practical protocol for long-term maintenance of gnotobiotic zebrafish and extend the utility of this model to study microbiome-dependent effects on host physiology, and development beyond early larval stages of life.

**IMPORTANCE:** While the gnotobiotic zebrafish have been a powerful model for interrogation of host-microbiota interactions, their use has been limited to early life stages due to complications of long-term husbandry. To address this limitation, we developed a simple protocol that enables rearing germ-free zebrafish well beyond larval stages. Germ free fish exhibit physiological and developmental defects that mirror those described in mammalian counterparts supporting a conserved role for microbiota in vertebrate development and physiology. Our protocol provides a method to investigate microbial influences on adaptive immunity, metabolism, and chronic disease processes in zebrafish not possible with current methodologies. Given the rapid and simple methods for gnotobiotic derivation and the large number of transgenic animal lines available for zebrafish we anticipate this model will accelerate mechanistic discovery of microbial impacts on host health.

## INTRODUCTION

The microbiome is the collection of microbes, their genes, metabolic products, and ecological conditions at a location^1,2,3^. Over the past 25 years our understanding of the beneficial functions of host associated microbiomes has substantially increased. We now know that the composition and functioning of microbial communities is a critical component of animal health^4–8^. For example, gut microbes help digest and absorb nutrients from our diets, produce essential micronutrients, regulate immune function and development, limit infections, and metabolize xenobiotics^9,5,10^. While disruption of these communities has been linked to diseases such as cancers^11^, obesity^12^, inflammatory bowel disease^13^, and neurodegenerative disorders^14^, the directionality of association and underlying molecular mechanisms are often poorly defined. These gaps in knowledge cloud the impact of these findings and complicate the development of microbiome modifying therapeutics.

Gnotobiotic animals were first developed in 1885 to explore host-microbe interactions and their impact on host health and disease^15,16^. Gnotobiotic mice have provided causal evidence that microbiota affects immunity^17^, brain function^18^, metabolism^19^, and development. However, the derivation of gnotobiotic mouse colonies is labor-intensive, time consuming, and costly. To complement the mouse, several low-cost gnotobiotic animal model systems have been developed and used to explore host-associated microbial ecology^20^. One such model, the gnotobiotic zebrafish, was developed in 2004^15^ as low-cost high-throughput model system for mechanistic investigations into host-microbiota interactions^21^. External fertilization and facile gnotobiotic derivation allow the generation of hundreds to thousands of gnotobiotic animals in less than a day. Growing use of gnotobiotic zebrafish has provided critical insights into microbial regulation nutrient metabolism and uptake^15,22^; pancreatic β-cell development^23^; intestinal epithelial proliferation^24^; and innate immune system development and function^25^. The transparent early life stages of fish enable real-time investigation of microbial community assembly and responses to various stimuli in the gut^26,27^. A major limitation of gnotobiotic zebrafish is that studies have been constrained to early life stages (<14 days post fertilization) due to reliance on live feeds (*e.g.*, *Tetrahymena*) for rearing young animals^15,28^. The lack of long-term gnotobiotic zebrafish husbandry protocols limits the utility of the model as 1) larval fish do not possess a functional adaptive immune system at this developmental stage^29,30^, 2) many organ systems are not fully developed this early in the zebrafish^31,32^, and 3) gut microbiomes are highly simplified at this stage compared to adult fish and mammals^33^.

The primary complication of long-term gnotobiotic culture in zebrafish is the lack of appropriate sterilized feeds. While gnotobiotic or UV sterilized live feeds^34,35^ have been used to extend the length of gnotobiotic zebrafish experiments, protocols for long term rearing of these animals (>14 days post fertilization) are lacking. Sterilization of live feeds is also laborious and increases the potential for contamination of gnotobiotic animals^36^. In mammals, autoclaved diets have been successfully used for gnotobiotic animal husbandry. However, consumption of autoclaved diets results in a progressive epidermal degeneration in zebrafish^15^. Another widely used method of sterilization of foods is gamma irradiation which damages DNA thereby inactivating microorganisms^40^. Currently, 40-50kGy doses of gamma irradiation are the gold standard for food sterilization for gnotobiotic mice^24^,however, it has not been investigated as a means for long-term culture of gnotobiotic zebrafish.

Here we investigated the feasibility of using dietary gamma irradiation as a method of long-term husbandry of germ-free zebrafish. In addition to its sterilizing effects, gamma irradiation can also damage dietary macro-and micronutrients leading to nutrient deficiencies, shifts in microbiome function, and immunomodulatory effects in the host^42,43^. So, we began by investigating the impact of feeding diets exposed to increasing doses of gamma-irradiated on zebrafish development and the microbiome structure. We then used these results to develop a long-term husbandry protocol for gnotobiotic zebrafish and evaluated the effects of microbiota and gnotobiotic derivation on gene expression and development. Our work demonstrates the feasibility of using gamma irradiation of dry diets for culture of gnotobiotic zebrafish far beyond previous standards and provides a foundational knowledge on the impact of the microbiome on zebrafish physiology.

## MATERIAL AND METHODS

### Animal experiments

*Animal husbandry.* Zebrafish (*Danio rerio* T5D strain) were housed on a 14:10 h light–dark cycle at 28°C and pH maintained between 7.0 - 7.3. Zebrafish were fed three times daily with GEMMA Micro 75 (Skretting, Westbrook, Maine) starting at five days post fertilization (dpf). Animal experiments were approved by the University of Illinois Institutional Animal Care and Use Committee.

*Experiment one - Evaluation of impact of gamma-irradiated diet consumption in microbiota replete animals*. Beginning at fertilization, zebrafish were housed in 75cm^2^ untreated polystyrene tissue culture flasks (VWR 10861-576) with five fish per flask, and six replicate flasks per group for 28 days. Fifty-percent water changes were performed every other day to maintain ammonia at or below 0.5ppm. This experiment was replicated twice (Replicate 1: N = 67; Replicate 2: N=70).

*Experiment two - Long-term gnotobiotic culture of zebrafish*. Germ-free (GF), conventionalized (CVZ) and conventional (CV) zebrafish were housed in eighteen 75cm^2^ flasks (six flasks per group; N=5 fish per flask). Germ-free animals were reared under sterile conditions after derivation; CVZ animals were derived gnotobiotic but were seeded with parental microbes immediately after derivation by exposure to parental tank water; CV animals were housed like GF and CVZ but did not undergo gnotobiotic derivation. An additional control group of fish from the same spawn as the CV, GF, and CVZ animals, colony controls (CC), were used to evaluate how consistent growth and physiology was between gnotobiotic animals and animals raised under standard husbandry conditions. Colony controls were housed in a 2.8L polycarbonate tank on a recirculating water system. The GF, CVZ, and CV groups were all fed a supplemented 20kGy irradiated diet, while the CC group were fed a unirradiated Gemma Micro 75 diet.

### Gnotobiotic derivation of zebrafish

Gnotobiotic T5D zebrafish were generated as described by Melancon et al. (2017)^44^. Embryos collected within one hour, were transferred to sterile antibiotic embryo medium (ABEM). Embryos were incubated at 28°C in ABEM for six hours, rinsed three times in sterile embryo media (EM), immersed in 0.1% povidone-iodine (PVP-I) for 2m, rinsed (3x) with sterile EM, immersed in 0.003% sodium hypochlorite for 20m, then rinsed (3x) with sterile EM. Sterilized embryos were transferred into 50mL sterile EM in 75cm^2^ flasks. Aliquots of EM visually inspected with phase-contrast microscopy and cultured twice weekly at 28°C on brain heart infusion agar to confirm sterility.

### Gamma irradiation of diets

Diets were sterilized by gamma-irradiation at different doses (10kGy, 20kGy, 40kGy and 80kGy) using a Gammacell Cobalt60 irradiator at the Nuclear Measurement Laboratory at the Department of Nuclear, Plasma, and Radiological Engineering, University of Illinois at Urbana-Champaign. The Gammacell 220^45^, manufactured by Nordion, is a self-contained Cobalt-60 irradiation unit consisting of sealed Co-60 sources enclosed within a lead shield, a cylindrical sample drawer, and a motorized drive mechanism that moves the drawer vertically along the source centerline. The irradiation cavity, measuring approximately 15 cm in diameter and 20 cm in height, can accommodate a range of sample types and sizes. The uniformity of the radiation field is ensured by the configuration of forty-eight double-sealed Co-60 source pencils, each 21.11 cm long, arranged in an annular formation around the chamber to provide a uniform gamma flux and minimize dose gradients. At the time of irradiation, the system delivered an average dose rate of approximately 2.1 kGy/h at the chamber center. The dose uniformity was validated using high-dose thermoluminescent dosimeters (TLDs) from Mirion, which confirmed an excellent homogeneity of the irradiation field along the chamber diameter, with a measured dose variation of only ±3% (±1 SD). In our second experiment, we fed GF, CVZ, and CV fish with 20kGy gamma-irradiated diets that had been supplemented with vitamins to mitigate potential loss of micronutrients impacted by irradiation or loss of vitamins synthesized by microbes. We supplemented irradiated Gemma Micro 75 with 5 mg Thiamine HCl, 10 mg Riboflavin, 10 mg Calcium pantothenate, 0.6 mg D-biotin, 4 mg Pyridoxine HCl, 1.5 mg Folic Acid, 200 mg Inositol, 60 mg L-vitamin C-2-Magnesium phosphate, 6.05 mg Niacin, 50 mg α-Vitamin E acetate, 4 mg Vitamin K, 0.11 mg Retinol acetate, 0.02 mg Vitamin D and 648.74 mg microcrystalline cellulose per gram feed (w/w) by spray-coating. Sterility of gamma-irradiated diets was confirmed with aerobic culture at 28°C for 24h on BHI agar plates.

### Amplicon sequencing

DNA was extracted from whole zebrafish using the DNeasy PowerSoil Kit (QIAGEN, Hilden, Germany) following the manufacturer’s protocol, with the addition of manual homogenization of whole larvae with a sterile pestle prior to the lysis. The V4 hypervariable region of the bacterial 16S rRNA gene was amplified from extracted DNA using the 515F and 806R barcoded primers^46^. Library concentration was quantified with the Qubit DNA HS assay kit (Life Technologies, Carlsbad, CA, USA) and 200 ng of each prepared library was pooled and cleaned with the QIAquick PCR purification kit (QIAGEN, Hilden, Germany). Libraries were then sequenced on an Illumina MiSeq at the Roy J. Carver Biotechnology Center at University of Illinois at Urbana-Champaign. This generated ∼10 million 300 bp paired-end reads in total (Experiment one: ∼5.3 million, Experiment two: ∼5 million)

### Microbial community analysis

*Experiment one*. Raw paired sequencing reads were filtered and trimmed using R (v4.5.0) and DADA2 (v1.20) with the following parameters: truncLen = c(240, 180), trimLeft = 7, maxEE = c(2,2), truncQ = 2, and rm.phix = TRUE. Forward and reverse reads were then denoised, merged, and subjected to chimera removal using the consensus method. Taxonomy was assigned using the SILVA (v138.1) reference database^47,48^.

*Experiment two*. Raw paired sequencing reads were filtered and trimmed using R (v4.5.0) and DADA2 (v1.20) with the following parameters: truncLen = c(250, 230), maxN = 0, maxEE = c(2,2), truncQ = 2, and rm.phix = TRUE. Reads were denoised, merged, chimera filtered, and assigned taxonomy as above.

*Diversity analysis and statistics*. Samples from experiment one and experiment two were rarefied to a depth of 7,000 and 70,000 reads, respectively. Samples with fewer reads than the rarefaction threshold were removed from subsequent analysis. Alpha diversity (richness and Shannon entropy) and beta diversity (jaccard) were calculated in R using the vegan package (v2.6-4). Principal component analysis (PCA) was performed using the stats package (v4.3.3) and visualized with ggplot2 (v3.5.0). Associations between microbiome beta diversity and feed and replicate parameters were quantified with permutational multivariate analysis of variance (vegan::adonis2, 5,000 random permutations, *P*<0.05). To assess the effects of experimental parameters on microbial diversity, generalized linear mixed models (GLMMs) were fitted using the glmmTMB package (v1.1.11).

### Transcriptome analysis

The AllPrep DNA/RNA Mini Kit (QIAGEN, Hilden, Germany) was used to isolate total RNA from whole zebrafish juveniles following the manufacturer’s protocol. Prior to lysis, GF, CVZ, CV, and CC samples were flash-frozen in liquid nitrogen and manually homogenized using a sterile pestle. The concentration of RNA was assessed with the Qubit RNA HS assay kit (Life Technologies, Carlsbad, CA, USA). RNA-Seq libraries were prepared with the Kapa Hyper Stranded mRNA library kit (Roche, Basel, Switzerland), pooled in equimolar concentrations, quantitated by qPCR, and sequenced on an Illumina NovaSeq X Plus (150 bp paired end).

FastQC (v0.11.9) assessed quality of the paired-end reads before and after trimming with Cutadapt v2.6 to remove adapters, low-quality reads (q<30), and reads shorter than 50bp, allowing a maximum of one ambiguous base (N). An additional 9 bases were trimmed from the 5’ end and 5 bases from the 3’ end of each read to remove low-quality bases and residual adaptor sequences^49^. STAR (v2.7.10a) aligned merged reads to the zebrafish reference genome (GRCz11; Ensembl^50^) using default parameters allowing a maximum of two mismatches and a minimum aligned read percentage of 30%^51^. Differential expression analysis was assessed with DESeq2 (v1.42.1)^52^ and a likelihood ratio test. Heatmaps were generated using the ComplexHeatmap (v2.18.0) R package^53^. Raw and normalized count plots were created via ggplot2 (v3.3.2). Gene set enrichment analysis (GSEA) was performed and visualized with gProfiler2 (v0.2.1)^54^ on differentially expressed genes (DEGs; adjusted p-value below 0.1, log2 fold change > 0.5) using six databases: Gene Ontology - Biological Process (GO:BP), Gene Ontology - Molecular Function (GO:MF), Kyoto Encyclopedia of Genes and Genomes (KEGG), Reactome, and TRANScription FACtor (TF). False discovery rate was controlled with the Benjamini‒Hochberg method (FDR < 0.2).

### Correlation analysis

Spearman’s rank correlation coefficient was calculated for all gene-bacteria taxa comparisons to assess the association between the variables. For each pair of variables, the Spearman correlation coefficient calculated in R (stats v4.5.0). The false discovery rate (FDR) was controlled (FDR < 0.1) using the Benjamini-Hochberg procedure.

### Data Availability

All sequencing data used in this manuscript is deposited at the National Center for Biotechnology Information Sequence Read Archive under project numbers PRJNA1338261 and PRJNA1338468 samples SAMN52403128-SAMN52403288 and SAMN52440818-SAMN52440835.

## RESULTS

### Gamma-irradiation alters microbiome diversity but not zebrafish growth

To determine if gamma irradiation of diets impacted zebrafish growth, we examined weight and length of animals fed diets dosed with 0-80kGy at 28dpf. As expected, gamma irradiation resulted in sterilization of feed at all doses examined while viable microbes were present in control feeds (**Supplemental Figure 1A**). Animals fed irradiated diets did not differ from controls in weight or length (Kruskall-Wallis; *P*>0.05, **Figure 1A, B**). Condition factor, an indicator of the fish health and growth similar to the body mass index^55^, was also unchanged between groups (*P*>0.05, **Figure 1C, Supplemental Table 1**). No differences in survival were noted between groups (Kruskal-Wallis; H=4.0, *P*=0.41). Together these results suggest that feeding gamma irradiated diets does not impact growth or survival of larval zebrafish.

**Figure 1:**
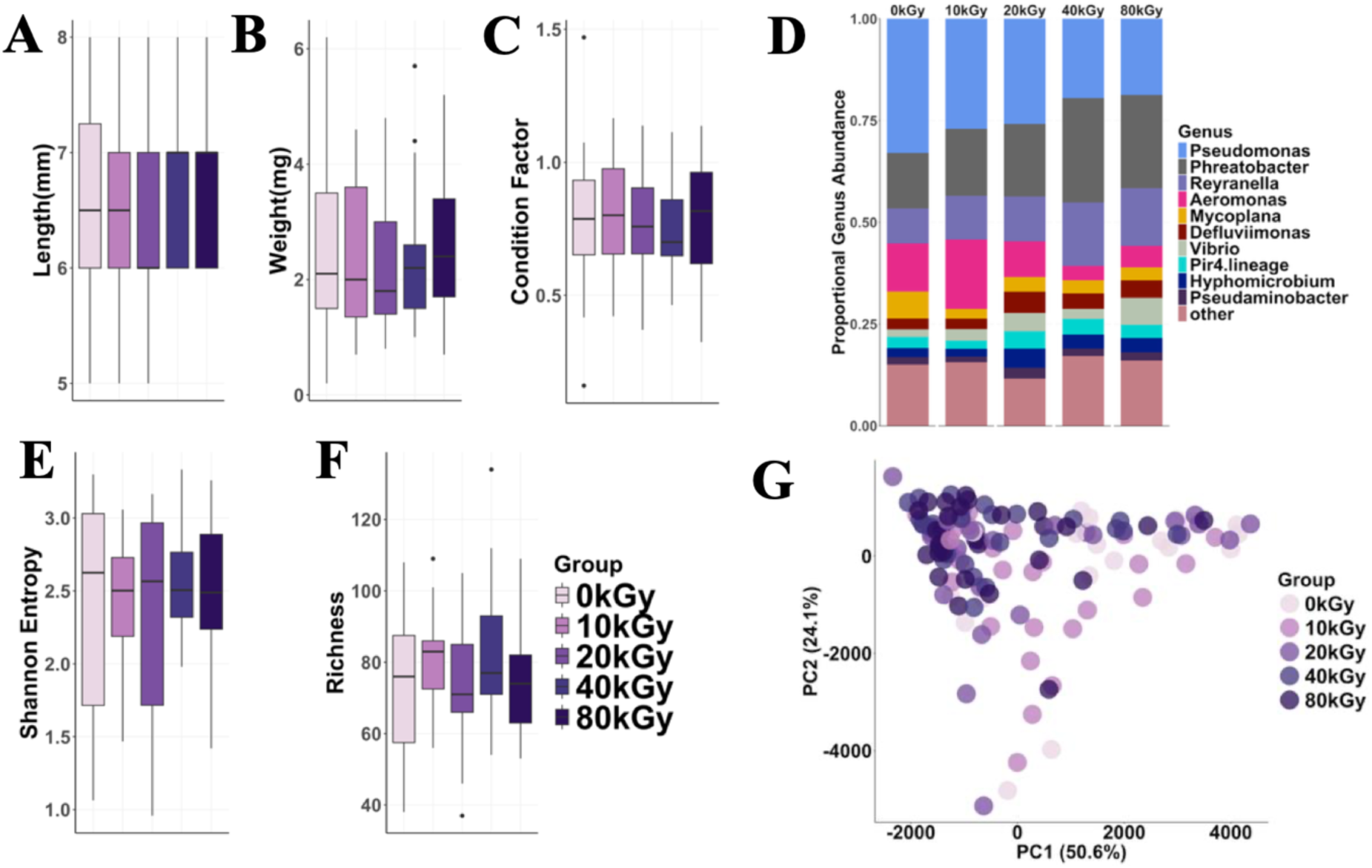
Gamma-irradiated diets do not affect zebrafish body metrics but alter microbiome composition. Boxplots show the effect of different diets on the **A)** Length, **B)** Weight, and **C)** Condition factor of zebrafish at 28 days post-fertilization (dpf). **D)** The relative abundance of the top ten microbial genera in the zebrafish. **E)** Shannon entropy and **F)** Microbial richness in zebrafish fed the various diets. **G)** Principal Component Analysis (PCA) ordination of microbial communities in zebrafish larvae fed gamma-irradiated diets compared to a non-irradiated diet (0 kGy, n = 27; 10 kGy, n = 27; 20 kGy, n = 29; 40 kGy, n = 25; 80 kGy, n = 29), with colors denoting specific diets.

Since the microbiome extensively links with health, we asked if consumption of irradiated diets alters microbiome diversity. The microbiota of animals fed irradiated and control diets were compositionally similar (**Figure 1D**). Neither microbial richness or Shannon entropy were significantly impacted by irradiation (Kruskal-Wallis; *P*>0.05; **Figure 1E-F**). However, community beta-diversity was associated with dose (PERMANOVA; R^2^=0.07, *P*=2.0×10^-4^; **Figure 1G**). To determine which taxa may be impacted by different diets, we fit negative binomial generalized linear mixed models (GLMMs) for each genus. We found 19 genera were significantly impacted by gamma irradiation including several highly abundant taxa (**Supplemental Figure 1B**; FDR<0.1). For example, *Phreatobacter* was significantly increased in the animals fed 40kGy (z=3.7, *P*=2.4×10^-4^) and 80kGy (z=2.9, *P*=1.2×10^-3^) diets compared to controls while *Stenotrophomonas* was elevated in 80kGy fed animals (z=3.3, *P*=1.1×10^-3^). Conversely, *Aeromonas*, abundance was decreased in 40kGy (z=-3.2, *P*=1.2×10^-3^) and 80kGy (z =-2.0, *P*=5.0×10^-2^).

One possible explanation for the changes in microbiome composition is that gamma irradiation killed microbes that normally would colonize the larval fish gut. If true, the compositions of microbiomes of control animals would be more similar to their diet than those fed irradiated diets. In contrast to the larval gut the feed microbiota was dominated by *Limosilactobacillus*, *Ligilactobacillus*, and *Candidatus Competibacter* (**Figure 1D, Supplemental Figure 1C**) and did not differ across dose (ANOSIM: R=0.11, *P*=0.30). Zebrafish microbiomes had significantly greater richness (Kruskal-Wallis; H=5.7, *P*=1.7×10^-2^) and lower Shannon entropy (Kruskal-Wallis; H=15.1, *P*=9.9×10^-5^) compared to diet (**Supplemental Figure 1D,E**). Microbial community diversity was associated with sample type (feed versus fish; PERMANOVA; R^2^=0.16*, P=*2×10^-4^; **Supplemental Figure 1F**) however, control diet microbiota were not more similar to their feed when compared to irradiated diet microbiota (*P* > 0.05, **Supplemental Figure 1G**). These data suggest that gamma-irradiated associated alterations zebrafish microbial communities are likely driven by factors other than reduction of viable biomass in diets.

### Gamma irradiation of diet enables long term husbandry of germ-free zebrafish

Next, we examined the feasibility of using gamma irradiated diets to rear germ-free zebrafish past the larval stage (∼2wk). We selected 8wk as a terminal time point as many aspects of social behaviors^56^, immune function^44^, and gastrointestinal system anatomy and function^57^, not yet fully matured at 2wk post fertilization, have been completed or advanced significantly at this time. We began by examining differences in growth in GF, CVZ, and conventional CV animals reared on 20 kGy diets for ∼8wks (55dpf) as well as CC reared under standard husbandry conditions (see methods). Weight (GLMM; CVZ: z=0.46, *P*=0.65; CV: z=2.74, *P*=6.0×10^-3^; CC: z=3.42 *P*=6.2×10^-4^) and length (CVZ: z=1.11, *P*=0.27; CV: z=3.65, *P*=2.7×10^-4^; CC: z=3.4, *P*=6.0×10^-4^) was significantly increased in CV and CC compared to GF, while the CVZ group did not differ significantly. Condition factor did not vary across group (*P*>0.05; **Figure 2A-C, Supplemental Table 2**).

**Figure 2:**
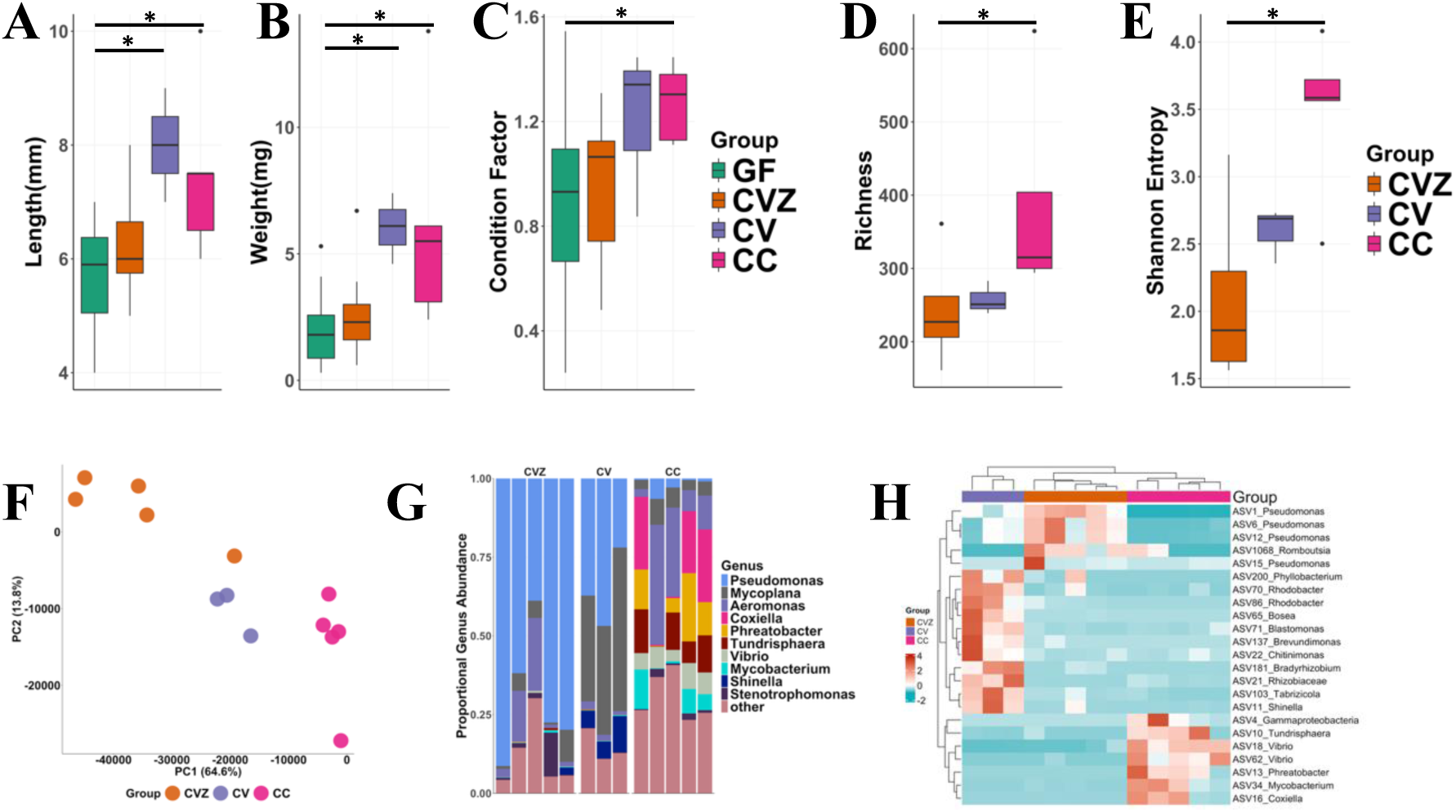
**Zebrafish growth metrics and microbiome composition differ across experimental groups**. Boxplots depict the **A)** length, **B)** weight, and **C)** condition factor of GF, CVZ, CV, and CC zebrafish at 55 days post-fertilization (dpf). **D)** Microbial richness and **E)** Shannon entropy boxplots for zebrafish in the CVZ, CV, and CC groups. **F)** Principal component analysis (PCA) ordination illustrating the clustering of microbial communities among CVZ, CV, and CC zebrafish, with colored dots representing different groups. **G)** Relative abundance of the top ten microbial genera across CVZ, CV, and CC zebrafish. **H)** Heatmap showing differentially abundant taxa among CVZ fish, CV fish, and CC fish. Each column represents an individual sample, and each row represents a taxon identified as significantly different among groups. Color intensity reflects Z-score normalized abundance across samples, with red indicating higher abundance and blue indicating lower abundance. Only taxa with p < 0.005 and q < 0.2 were included.

Since husbandry and derivation conditions differed between groups, we examined how microbial diversity and composition differed across groups. Richness was increased in CV (z=6.16, *P*=0.83) and CC animals (CC: z=2.58, *P*=1.0×10^-2^) compared to CVZ (**Figure 2D**). Shannon entropy increased only in CC animals (*P*=0.04; **Figure 2E**). Community beta diversity also associated with group (R^2^=0.48, *P*=4×10^-4^), (**Figure 2F**). Consistently, differences were also noted in genera (**Figure 2G, Supplemental Table 3**) and ASV abundances (**Figure 2H**, **Supplemental Table 4**). Highly abundant ASVs corresponding to the genus *Pseudomonas* (ASV1, ASV6, ASV12) and *Shinella* (ASV11) where significantly elevated in CVZ fish compared to CC animals. ASVs associated with the genus *Phreatobacter* (ASV13, ASV66), *Vibrio* (ASV18), and *Coxiella* (ASV16) were depleted in CVZ and CV (**Supplemental Table 4**). Together these data indicate that gnotobiotic derivation or gamma irradiation likely influence microbial community structure.

### Germ free animals manifest unique gene expression profiles

To investigate impact of microbial communities on host physiology, we conducted whole animal RNA-Seq of individual GF, CVZ, CV, and CC zebrafish. In total, 2,615 DEGs (FDR=0.1) were detected between groups (**Figure 3A, Supplemental Table 5**) with group explaining ∼27% of variance in gene expression (R^2^=0.27, *P*=6.0×10^-3^; **Figure 3B**). When compared to GF, 1,441; 2,268; and 2,500 genes were differentially expressed, respectively (**Figure 3C**). Of these, 1,080 genes showed consistent directional changes across these groups. An additional 973 genes were consistently differentially regulated when CC and CV were compared GF animals and a further 322 were uniquely different between CC and GF.

**Figure 3:**
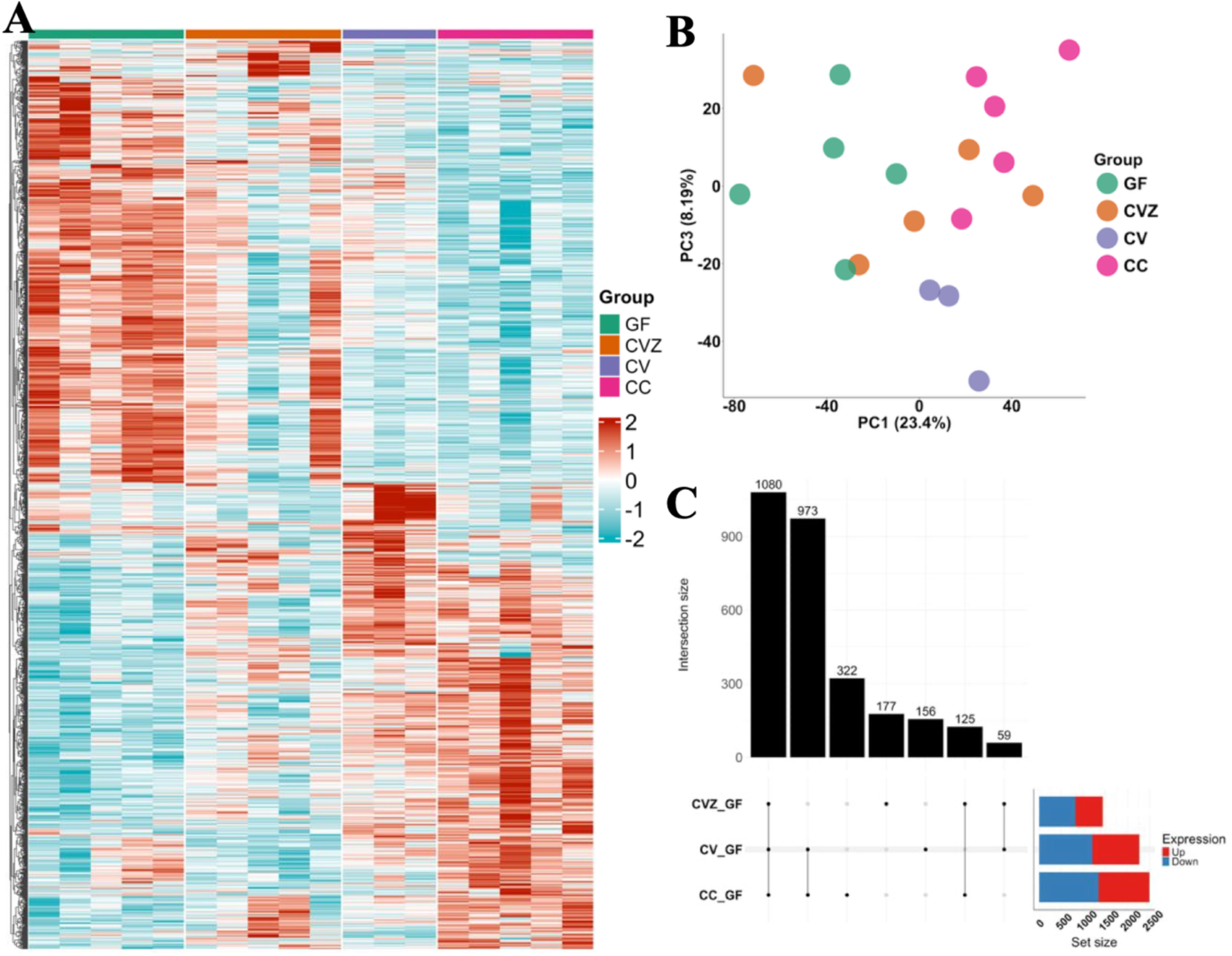
Germ-free zebrafish exhibit distinct gene expression profiles compared to microbe colonized animals. **A)** Heatmap showing the differentially expressed genes in GF, CVZ, CV and CC fish. The different groups are indicated by colored bars on the top. **B)** Principal Component analysis (PCA) plot showing the clustering of expressed genes in GF, CVZ, CV, and CC fish **C)** Upset plot showing the intersection of shared and unique genes of CV, CVZ and CC fish in reference to germ-free fish (GF)

To gain deeper insights into how these differences gene expression may impact host physiology, we evaluated gene set enrichment for differentially expressed genes. In total, 763 terms (GO:BP = 522, GO:MF = 194, KEGG = 11, REAC = 29, TF = 7) were significantly enriched (FDR ≤ 0.2) for up-and down-regulated genes in GF animals compared to controls (**Figure 4, Supplemental Table 6**). Consistent with previous work in germ-free zebrafish^58^, terms involved in microbial sensing and immune responses including response to lipopolysaccharide (GO:0032496), innate immune system (R-DRE-168249), neutrophil degranulation (R-DRE-6798695) and T cell costimulation (GO:0031295) were enriched for genes downregulated in GF zebrafish. Pathways involved in maintenance of mucosal barrier integrity including tight junction assembly (GO:0120192) were also enriched in downregulated GF genes. Nutrient uptake, metabolism, and deposition were also significantly altered in germ-free animals. Lipid localization (GO:0010876), binding (GO:0008289), transport (GO:0006869), and metabolic process (GO:0006629) were all enriched for genes downregulated in GF animals. Notably, PPAR signaling pathway (KEGG:ko03320), which plays an important role in lipid metabolism and adipocyte development, was also enriched for genes downregulated in GF animals. Upregulated genes in GF animals also enriched for pathways associated with growth (*e.g.,* developmental growth – GO:0048589 and growth - GO:0040007), cell death (programmed cell death - GO:0012501), and development (*e.g.,* eye development - GO:0001654; central nervous system development - GO:0007417; neuron differentiation - GO:0030182).

**Figure 4.**
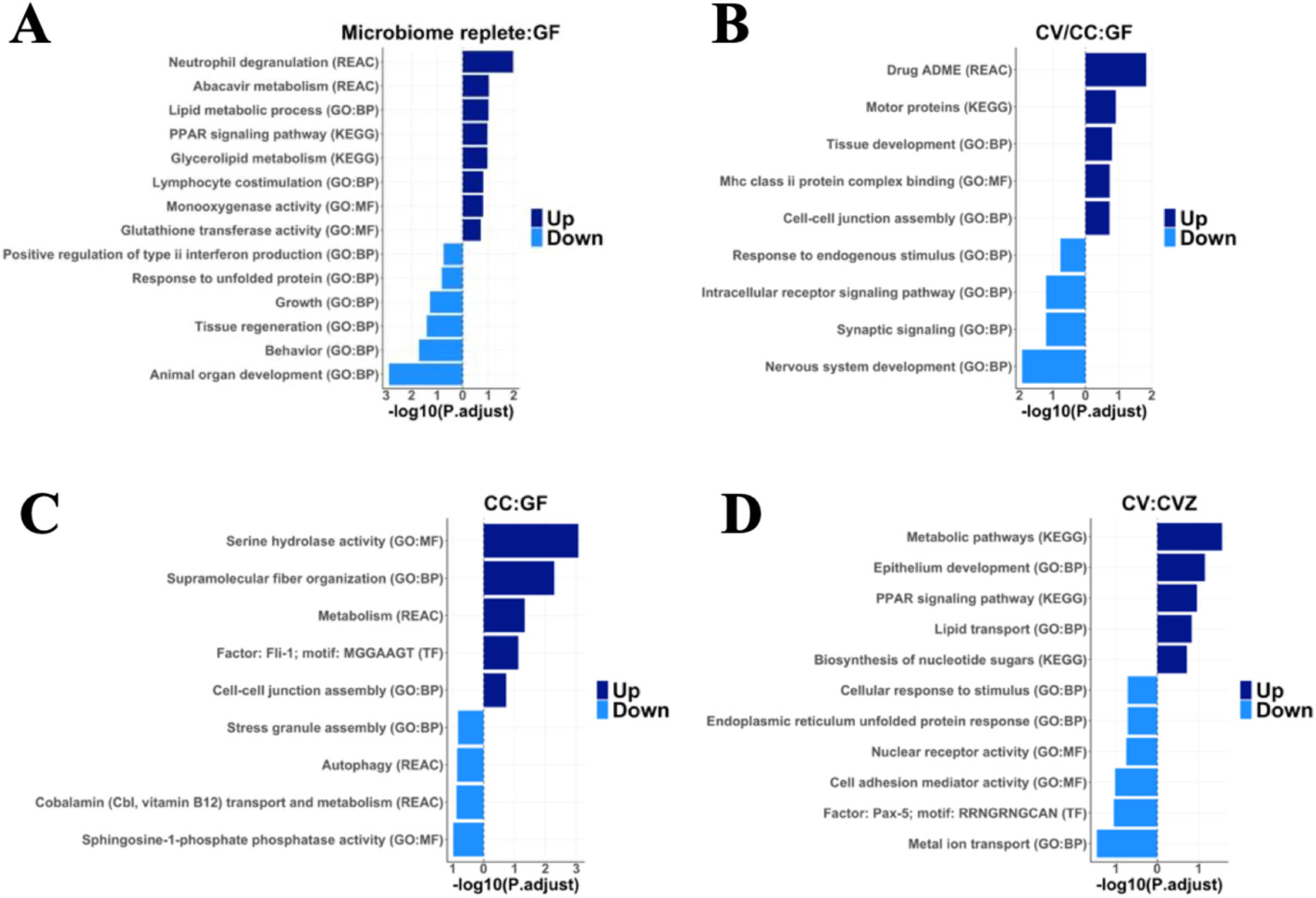
Enriched pathways associated with host gene regulation in response to the microbiome. (A-B) Selected pathways enriched among genes regulated in the same direction in **A)** microbiome-replete versus germ-free (GF) animals and **B)** CV and CC animals versus GF. (C-D) Selected pathways enriched among differentially expressed genes in **C)** CC versus GF animals and **D)** CV versus CVZ animals. Bars represent the significance of enrichment (-log10 adjusted P-values). Pathways associated with upregulated genes are indicated in darker blue, while those associated with downregulated genes are shown in lighter blue. The dashed vertical line denotes the baseline for no enrichment. Database sources are indicated in parentheses: Gene Ontology - Biological Process (GO:BP), Gene Ontology - Molecular Function (GO:MF), Kyoto Encyclopedia of Genes and Genomes (KEGG), the Reactome database (REAC), and TRANScription FACtor (TF) database.

Animals that did not undergo GF derivation (CC and CV) uniquely shared up-regulated genes enriched in pathways involved in drug absorption, distribution, metabolism, and excretion (ADME) (REAC:R-DRE-9748784), immune function (GO:0007043, GO:0030199, GO:0023026, GO:0042605), and lipid metabolism (GO:0042577; **Figure 4B**). Genes uniquely down-regulated in CC compared to GF were enriched in pathways such as autophagy (REAC:R-DRE-9612973, REAC:R-DRE-1632852), stress granule assembly (GO:0034063), vitamin B12 metabolism (REAC:R-DRE-196741, **Figure 4C**) while upregulated genes were enriched in drug ADME (REAC:R-DRE-9748784), and metabolism (REAC:R-DRE-1430728, KEGG:ko00500.

Comparing CVZ to CV fish (**Figure 4D**), we observed that pathways including epithelium development (GO:0060429), PPAR signaling (KEGG:ko03320), metabolic pathways (KEGG:ko01100), lipid transport (GO:0006869) were enriched for genes upregulated in CV animals. Pathways such as endoplasmic reticulum unfolded protein response (GO:0030968), Factor: Pax-5; motif: RRNGRNGCAN (TF:M03577), metal ion transport (GO:0030001), cellular response to stimulus (GO:0051716), cell adhesion mediator activity (GO:0098631) were enriched CV upregulated genes (**Figure 4D**).

### Microbiome abundances correlate with host transcriptome profiles

To determine if altered gene expression associated with differences in microbial community composition, we calculated Spearman correlation coefficients between all differentially abundant ASVs and DEGs. After filtering modest correlations, we found 2,555 associations (|ρ|>0.5, FDR<0.1) between 24 unique taxa and 1,126 genes (**Figure 5, Supplemental Table 7**). Many of these associations involved abundant commensals of zebrafish larvae including *Pseudomonas* (837), *Vibrio* (285), and *Phreatobacter* (207). These taxa often exhibit patterns of association with host genes that are the inverse of each other. For example, several *Pseudomonas* ASVs negatively correlated with several genes involved in immune function and cell death (**Figure 5A**) and lipid metabolic processes (**Figure 5B**) while *Vibrio* ASV show the opposite trend. Like *Vibrio* several ASVs belonging to *Tundrisphaera*, *Phreatobacter*, and *Mycobacterium* were positively correlated with critical genes involved in lipid metabolism and uptake including *gstt1a*, *fabp1p.1*, *cyp2p9, cd36*. Expression of several key genes involved in immune functioning and apoptosis (*casp22*, *trim35-3*, *ncf1*, *lyn*, *stat4*) were all also linked to these and other taxa. Similar patterns were seen with microbial associations with genes involved in response to xenobiotics (**Figure 5C**) with key regulators of these processes (*cyp2×9*, *cyp2×8*, *cyp2p*, and *gstt1a*) exhibiting distinct patterns of association with *Pseudomonas*, *Vibrio*, *Phreatobacter* ASVs. Finally, growth, development, differentiation including several involved in Notch and Wnt signaling pathways were linked to numerous taxa (**Figure 5D**). Collectively, these data support the hypothesis that the presence of a diverse microbiota influence host metabolism, immune stimulation, and response to xenobiotics.

**Figure 5:**
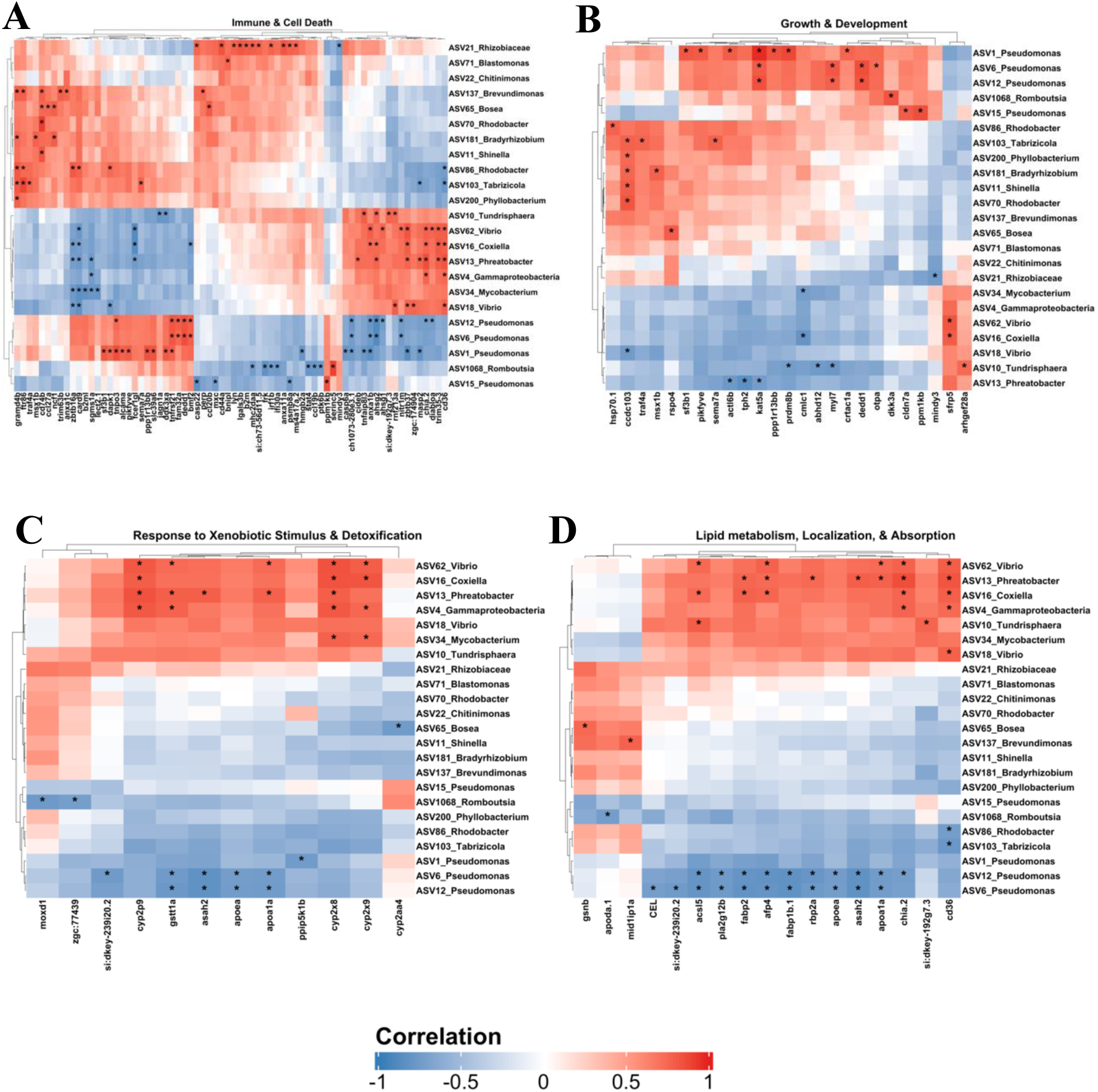
Correlation heatmap of a subset of differentially regulated genes and bacterial taxa. The heatmap illustrates the significant correlations between differentially expressed genes (columns) and bacterial taxa (rows). Colors represent the strength and direction of correlations, with red indicating positive correlations and blue indicating negative correlations. Statistically significant correlations (FDR< 0.1, |ρ| > 0.5) are shown with an asterisk. The side dendograms illustrate hierarchical clustering of genes and taxa. Panels correspond to gene categories: **A)** Immune & Cell Death, **B)** Growth & Development, **C)** Response to Xenobiotic Stimulus & Detoxification and **D)** Lipid Metabolism, Localization & Absorption.

## DISCUSSION

Host associated microbiota play an important role in development^59,60^, immune function^61–64^, metabolism^65,66^, and behavior^67,68^. Gnotobiotic animals offer a tool to explore the mechanistic underpinnings of these associations. The simple methods for gnotobiotic derivation, amenability to high-throughput screening, and transparent early life stages of the zebrafish has led to growing interest in this model. However, constraints on the experimental length and life stages available for investigation has limited its potential. Our work shows that gamma irradiation of pelleted feeds provides a viable method of rearing gnotobiotic zebrafish beyond early larval stages of development. Our approach eliminates barriers for adoption of this model system and has the potential to fast-track future mechanistic studies into host-microbe interactions across developmental stages.

Any long-term method for rearing gnotobiotic zebrafish should minimize impacts on development^35^. The impact of irradiated diets was minimal in microbiota replete animals with only modest differences in microbial community composition. More significant impacts on microbial community composition were observed between GF control groups (CC, CV, and CVZ), despite all groups receiving the same exposure to parental microbiota. While modest changes in microbiome composition in microbiota replete animals may be due to stochastic effects or destruction of nutrients caused by irradiation^41,42^, the causes for the more substantial effects in GF controls could have several causes. For example, CC exhibited the most diverse microbiota potentially due to housing and diet differences which are known determinants of diversity in zebrafish and other animal models^69,70^. Previous work has also shown that timing of exposure, microbiome diversity, and initial inoculum composition impact microbial assembly and host development^71–73^. It is also possible that residual antimicrobials or disinfectants used in derivation impacted microbial colonization and succession in the CV and CVZ animals^74,75^. Since our data suggest that the resulting communities associate with host gene expression, future work should attempt to minimize the compositional differences between GF experimental controls (*i.e.*, CVZ and CV) and animals reared under standard conditions.

While early larval stages rely primarily on yolk reserves^76^ and show minimal size differences, growth deficits become evident in our model as zebrafish have transitioned to exogenous feeding. Specifically, juvenile GF zebrafish exhibited reduced body size and weight. Similar observations have been made in other gnotobiotic animal models, including pigs and mice^77,78^. Germ-free piglets exhibit impaired growth, and nutrient digestibility a phenotype that can be reversed with microbiota transplantation^77^. It is well established that host associated microbiota play a critical role in host nutrient absorption and metabolism^4,79^ so it follows that the reduced weight in our GF animals may reflect defects in these pathways^22,80^. However, a major limitation of current GF zebrafish models is that animals are not fed making investigations of microbial interactions with host nutrient uptake and metabolism impossible^81^. Our protocol enables these investigations and our results mirror those in mammalian systems. For example, GF zebrafish fed irradiated diets have reduced expression of intestinal lipid transporters and fatty acid–binding proteins in the absence of microbes^6,8,58^. The decrease in lipid metabolism and localization is also consistent with reports in antibiotic treated zebrafish^82^. We also observed that genes involved in bile acid homeostasis including *akr1b1.2*, *ugt2a6*, *apoa1a*, *cyp27a2*, and *cyp3a65*, were downregulated in GF fish compared to microbe colonized fish, consistent with microbial modulation of bile acid metabolism in other vertebrates^83,84^. We also find that many of these differences link to specific microbiota including abundant commensals of the zebrafish gut. Specifically, abundance of ASVs associated with the genus *Pseudomonas* were negatively correlated with expression of *apoa1a*, *fabp2*, *fabp1b.1*, and *acsl5*, indicating the possible involvement of these taxa in regulation of fatty acid transport and activation pathways. Consistently, marked increased abundance of *Pseudomonas* was noted in the smaller CVZ and CV animals compared to the larger CC controls. Whether these observations are cause or consequence of differences in lipid metabolism and the mechanisms that underpin these associations is unclear and warrants further investigation.

Zebrafish are an important model system for liver disease and toxicology^85^ as well as the role the microbiome plays in these processes^27,86^. We found that our GF animals had significantly lower expression liver development and hepatocyte differentiation genes. Juvenile GF fish also had decreased levels of genes involved in xenobiotic metabolism and sensing in the GF zebrafish This finding aligns with the prior work suggesting that the GF zebrafish may be less capable of detoxifying dietary constituents and environmental components^15^. These shifts support a role for microbes in promoting liver development, function, detoxification as demonstrated in other models^87^. Our model now provides opportunities for reductionist study gut-liver-microbiota interactions and their impact on chronic disease processes and detoxification.

Reciprocal relationships between them microbiome and host are crucial for development, training of innate and adaptive immune systems and maintenance of their interactions with microbiota^88,29^. Zebrafish possess a functional innate immune system within days of fertilization, while their adaptive immune system becomes mature and functional within 4 - 6 weeks of hatch^89^. Germ-free animals exhibited numerous developmental impairments, including diminished cellular components of mucosal and systemic immunity^88,90–93^. Consistently, GF juvenile fish had decreased expression of genes involved in lymphocyte proliferation and innate and adaptive immune responses. One limitation of this work was that we use bulk RNA-Seq of whole animals, which likely limited our ability to resolve gene expression heterogeneity particularly in minority cell populations. Application of single cell/nuclei or spatial transcriptomic sequencing could help resolve more precisely differences in immune cell subsets in GF animals^94^. Such work would provide whole animal resolved investigations of microbial impacts on innate and adaptive immune development, response, and homeostasis.

Our study presents a significant advance in the field of zebrafish gnotobiotic research with the potential to enable low-cost mechanistic study of microbiota interactions with their hosts. One limitation of this work is that animals were only raised until juvenile phases of development. Ongoing experiments in our lab seek to rear and spawn adult germ-free zebrafish for the first time. Slower growth and developmental differences, while common in other gnotobiotic models, should also be addressed with future work that defines the optimal nutritional parameters, and housing conditions for gnotobiotic and conventionally raised zebrafish. Despite these caveats, our approach provides a foundation for future work that can leverage this system to define mechanisms of interactions between host and microbiota across lifespan.

## Acknowledgments

Research reported in this publication was supported by the National Institute of Environmental Health Sciences of the National Institutes of Health under Award Number R01ES036174 to CAG. The content is solely the responsibility of the authors and does not necessarily represent the official views of the National Institutes of Health. This work was also supported in part by a Cancer Center at Illinois pilot award, and UIUC start-up funds to CAG. The authors would like to thank the University of Illinois Urbana-Champaign (UIUC) DNA Services Core at the Roy J. Carver Biotechnology Center for their assistance with sequencing. This work made use of the equipment, software, and facilities provided by the University of Illinois Urbana Champaign College of Veterinary Medicine Shared Equipment Program’s Biocomputing Shared Resource (BioShaRe). The College of Veterinary Medicine BioShaRe is housed in the Illinois Campus Cluster, a computing resource that is operated by the Illinois Campus Cluster Program (ICCP) in conjunction with the National Center for Supercomputing Applications (NCSA) and which is supported by funds from the University of Illinois at Urbana-Champaign.

