## Supplementary figures and images for "Long Term Culture of Germ-Free Zebrafish Using Gamma-Irradiated Feeds"

### supplemental figure 1

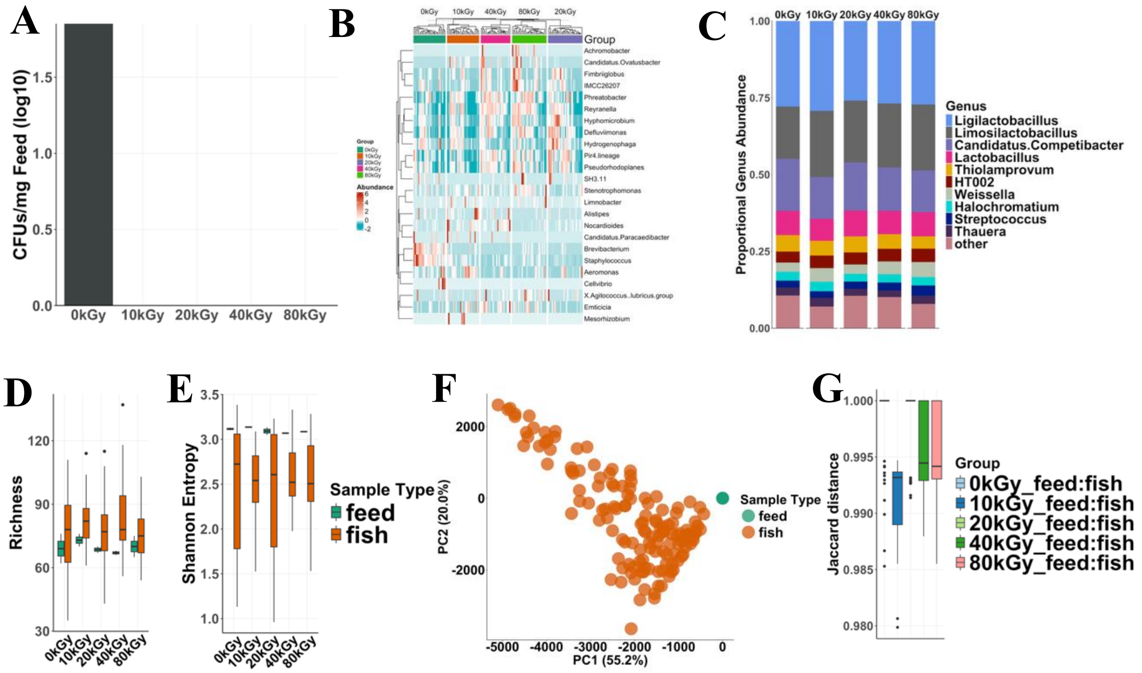


#### Supplemental Figure 1
